# New Drosophila long-term memory genes revealed by assessing computational function prediction methods

**DOI:** 10.1101/414565

**Authors:** Balint Z. Kacsoh, Stephen Barton, Yuxiang Jiang, Naihui Zhou, Sean D. Mooney, Iddo Friedberg, Predrag Radivojac, Casey S. Greene, Giovanni Bosco

## Abstract

A major bottleneck to our understanding of the genetic and molecular foundation of life lies in the ability to assign function to a gene and, subsequently, a protein. Traditional molecular and genetic experiments can provide the most reliable forms of identification, but are generally low-throughput, making such discovery and assignment a daunting task. The bottleneck has led to an increasing role for computational approaches. The Critical Assessment of Functional Annotation (CAFA) effort seeks to measure the performance of computational methods. In CAFA3 we performed selected screens, including an effort focused on long-term memory. We used homology and previous CAFA predictions to identify 29 key *Drosophila* genes, which we tested via a long-term memory screen. We identify 11 novel genes that are involved in long-term memory formation and show a high level of connectivity with previously identified learning and memory genes. Our study provides first higher-order behavioral assay and organism screen used for CAFA assessments and revealed previously uncharacterized roles of multiple genes as possible regulators of neuronal plasticity at the boundary of information acquisition and memory formation.

## INTRODUCTION

In the sequencing age, public data present the opportunity to the experimental biologist to address important biological questions that might otherwise remain hidden(Faith *et al.* 2007, e8; Yan *et al.* 2010, e12139; Chikina *et al.* 2009, e1000417; Chikina and Troyanskaya 2011, e1001074). However, the data abundance and availability can be a daunting project to tackle. Thus, we ask the question, how and to what extent can one utilize so much data to ask a very particular series of questions?

Data-mining algorithms have been designed and subsequently utilized to dissect large data sets. This utilization can present the discovery of novel and biologically relevant pathways and gene functions that might be overlooked due to the inability to pass detection threshold of other methodologies(Greene *et al.* 2009, 1289-1296; Greene *et al.* 2014, 1896-1900; Greene and Troyanskaya 2012, 95-100; Greene and Troyanskaya 2011, W368-74). By utilizing a bioinformatics-based approach, an accurate computer-generated list of gene targets can be generated to shed light on important biological process. This allows the biologist to rapidly conceive of, and subsequently test, new hypotheses on new targets that would be otherwise overlooked. Evaluating such methods frequently relies on curated databases of gene functions derived from the literature. We sought to complement these evaluations with assays of the behavioral phenomenon: long-term memory formation.

The ability to perceive, process, respond to, and remember past, present, and potentially upcoming environmental challenges is essential for survival. This ability is observed across the animal kingdom and remains a ripe, and understudied field of study. The field of memory and the accompanying neurobiology has arisen from the utilization of multiple approaches, ranging from behavior testing, to experimental perturbation that includes pharmacological, biochemical, and anatomical stressors in model and non-model systems(Dubnau *et al.* 2001a, 476; Dubnau and Tully 1998, 407-444; Dubnau *et al.* 2001b, 476-480; Margulies *et al.* 2005, R700-R713; Tully *et al.* 1994a, 35-47; Tully 1987, 330-335). Genetic manipulations have provided key insight into behavioral plasticity that facilitates memory formation and retention. The use of genetics allows the researcher to understand the hereditary components that regulate neuronal plasticity. Use of genetic systems allows one to find clues illuminating the extent of evolutionary conservation among the different molecular mechanisms that govern learning and memory (Greenspan 1995, 747-750).

*Drosophila melanogaster* has been utilized as a model system to understand the genetic and molecular basis of memory. Using associative(McGuire *et al.* 2005, 328-347; Tully *et al.* 1994b, 35-47; Tully 1987, 330-335; Tully and Quinn 1985, 263-277) and non-associative paradigms(Das *et al.* 2011, E646-54; McCann *et al.* 2011, E655-62; Ramaswami *et al.* 2013, 727-736; Ramaswami 2014, 1216-1229), an insight into the molecular mechanism governing the formation of long-term memories has been provided. However, the genes and proteins discovered in this manner might only be a small representation of the underlying learning and memory factors (i.e. genes, pathways, stimuli) that manifest in the insect brain. Therefore, it remains valuable to develop and utilize ways in which we may define potentially new genes and proteins. These novel elements that may be involved in the complex process of memory can be identified through an unbiased approach and subsequent mechanistic analysis.

There exists a strong need to have a means to understand the process of genes and gene function, including within the study of memory. The approach to understanding this process can be and has been expedited through the use of computational algorithms. These algorithms aim to automatically annotate and categorize genes to a given function. Genetic annotation via these algorithms can be based on multiple factors including the nucleic acid sequence and similarity to other known genetic elements (Wilson *et al.* 2000, 233-249; Cozzetto and Jones 2017, 55-67), evolutionary relationships(Pellegrini *et al.* 1999, 4285-4288; Engelhardt *et al.* 2005, e45), the gene expression patterns(Greene and Troyanskaya 2011, W368-74; Greene *et al.* 2014, 1896-1900; Greene *et al.* 2016, 557-567; Greene and Troyanskaya 2012, 95-100), the gene interaction partner(s)(Marcotte *et al.* 1999, 751-753; Vazquez *et al.* 2003, 697), and other features(Lan *et al.* 2013, S8; Sokolov *et al.* 2013, S10; Cozzetto *et al.* 2016, 31865). Given so many possible modes of prediction, there exists a need for a commonly accepted and utilized modality. The <u>c</u>ritical <u>a</u>ssessment of <u>f</u>unctional <u>a</u>nnotation (CAFA) was conducted to address this need (Jiang *et al.* 2016,184; Radivojac *et al.* 2013, 221). CAFA has historically relied on curated databases of gene functions, but there remains a need to complement these literature-derived resources with biological assays(Greene *et al.* 2016, 557-567). This remains especially true in biological processes, such as learning and memory, where relatively few proteins have experimental annotations.

The inherent plasticity present in an organism allowing it to recall information can allow the individuals and/or groups to modify many facets of their behavior, including reproduction. Modulation of reproductive behavior, and anticipation of conditions that effect offspring survival, which in turn affects fitness, is necessary for the survival of an organism. For example, *Drosophila melanogaster* females will retain their eggs (oviposition depression) when environmental conditions are determined to be nutritionally unsuitable(Spradling *et al.* 1999, 135-177). Drosophila will also depress their oviposition rates during and following an encounter with an endoparasitoid wasp by inducing effector caspases in a stage specific manner in the ovary(Lefevre *et al.* 2012, 230-233; Lynch *et al.* 2016; Kacsoh *et al.* 2017; Kacsoh *et al.* 2018, e1007430; Kacsoh *et al.* 2015c, 10.7554/eLife.07423). These endoparasitoids regularly infect Drosophila larvae, upwards of 90% in nature(Driessen *et al.* 1989, 409-427; Fleury *et al.* 2004, 181-194; LaSalle 1993, 197-215). Egg depression following wasp-exposure is maintained and is dependent on long-term memory formation(Kacsoh *et al.* 2018, e1007430; Kacsoh *et al.* 2017; Kacsoh *et al.* 2015c, 10.7554/eLife.07423; Bozler *et al.* 2017, e1007054). It is important to note that adult Drosophila are under no threat from these wasps. This makes all behavioral changes in response to the wasps anticipatory as a means to benefit their offspring. These changes are mediated, in part, by the visual system (Kacsoh *et al.* 2018, e1007430; Kacsoh *et al.* 2015c, 10.7554/eLife.07423; Kacsoh *et al.* 2015a, 1143-1157; Kacsoh *et al.* 2013, 947-950). There has also been evidence to demonstrate the involvement of long-term memory genes and proteins that mediate the maintenance of this oviposition depression behavior. In particular, the role of these genes in the <u>m</u>ushroom <u>b</u>ody (MB), the region of the fly brain implicated in learning and memory(Aso *et al.* 2009, 156-172; Aso *et al.* 2014, e04577; Claridge-Chang *et al.* 2009, 405-415; Masse *et al.* 2009, R700-R713; Schwaerzel *et al.* 2003, 10495-10502), has been demonstrated to function in the maintenance of the egg-depression behavior(Kacsoh *et al.* 2017; Kacsoh *et al.* 2015c, 10.7554/eLife.07423; Kacsoh *et al.* 2015a, 1143-1157; Kacsoh *et al.* 2018, e1007430; Kacsoh *et al.* 2015b, 1143-1157; Bozler *et al.* 2017, e1007054).

Given the current findings, we might only be scratching the surface of the alien world that is the foundation of neurobiology and the study of memory—not just in the Drosophila brain, but other organisms. These yet-to-be-identified factors might be necessary and/or sufficient for the processes of information acquisition, memory induction and maintenance, following the ability of memory recall. Given this extremely multifaceted process containing multiple layers of signaling, it remains valuable to identify and test novel genetic candidates. In order to find new genes and pathways, we must identify, create, implement, and validate new methodologies, which are independent of classical mutagenesis-based approaches.

In this study, we performed assays for CAFA3 designed to assess the performance of algorithms predicting novel genetic candidates that may be involved in long-term memory. We selected 29 genes to screen that we expected would be particularly useful for this evaluation. This manuscript focuses on the biological findings resulting from the screen. Through the use of RNA interference (RNAi) of each gene by expression of an RNAi transgene in the MB, we identified 3 genes involved in perception of the stimulus (wasp exposure), and 12 genes involved in long-term memory formation. Using a functional relationship network analysis, we find that the 3 perception genes are enriched in developmental and response to stimuli processes, while our long-term memory candidates are highly connected to learning and memory genes, including ones previously identified to be involved in this particular behavioral paradigm, and that the network is enriched for the process of long-term memory.

## RESULTS

### CAFA ANALYSIS REVEALS NOVEL GENE TARGETS

In order to identify novel genes involved in learning and memory and evaluate prediction tools, we turned to a computational approach using the principles of active learning(Sverchkov and Craven 2017, e1005466). Twenty-nine candidate genes for screening were selected based on an ensemble of top performing predictors in the CAFA2 challenge (Jiang *et al.* 2016, 184). Specifically, we collected prediction scores from top ten computational models that submitted predictions on 3195 Drosophila target genes (Figure 1 A). Their raw prediction scores on each gene for the biological process “long-term memory” (GO:0007616) were first converted into quartiles (*Q*) with five possible outcomes from 0 indicating a negative prediction up to 1 indicating a positive prediction with a step size 0.25. We then averaged the quartiles from these ten models for each gene so as to group them into three categories: (i) “likely positive” with average *Q* greater than or equal to 0.5 and covered by at least 7 models; (ii) “likely negative” with average *Q* of 0.00 and agreed by at least 7 models; and (ii) “variable” with average *Q* between 0.1 and 0.25 and standard deviation greater than 90% quantile (i.e., top 10% variable) and covered by at least 7 models (Figure 1 B). After excluding essential genes and those known to be associated with the GO:0007616 term by FlyBase, we randomly selected subsets of genes from the positive class (Mob4, rut, CG17119, GluRIA, Snap25), the negative class (Rnp4F, sens, sna, Capr, CG12744, CG5815, Oaz, Trpm, wdn) and the variable class (5-HT7, Cad87A, Dop1R1, Dop1R2, mnb, Nrk, Octbeta1R, CG18812, ft, knurl, p38c, sir, sdk, shf, TkR86C).

**Figure 1.**
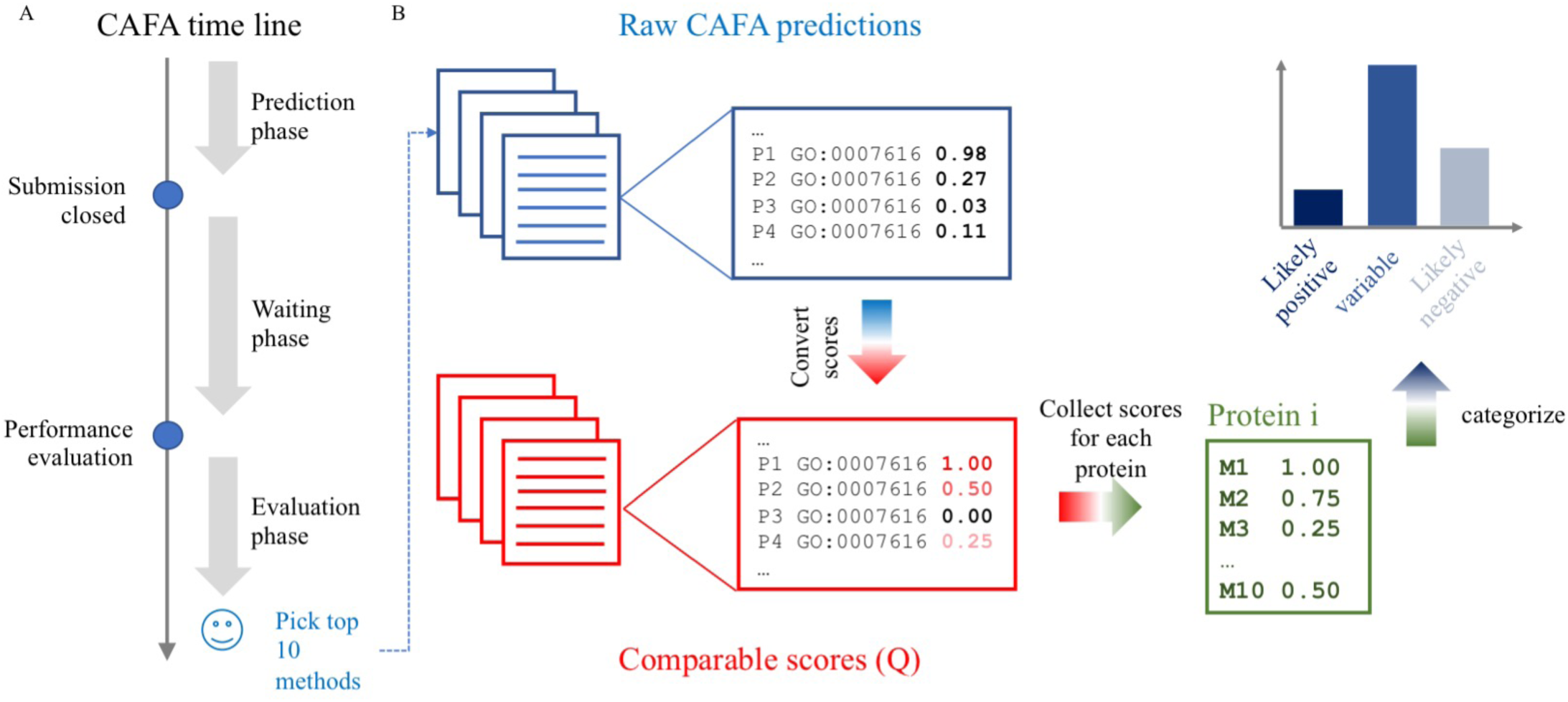
CAFA3 pipeline utilized in this study. Experimental design for CAFA3 algorithm (A). (A) illustrates the time line for CAFA challenge where we collect top-performing methods for Drosophila genes on GO:0007616. (B) indicates the process where we convert raw prediction scores to quantile (Q) and accumulate for each protein. We subsequently group these genes into three categories, where we picked 29 candidates for experimental verification.

### FLIES DEPRESS OVIPOSITION DURING AND FOLLOWING WASP EXPOSURE

In order to elucidate the roles of these genes in memory, we turned to an assay using the Drosophila-endoparasitoid-wasp system. Our assay utilizes the behavior of *D. melanogaster* following exposure to parasitoid wasps. One of the behaviors during and following wasp encounter is manifested by female flies that depress their egg-laying (oviposition) rate during wasp exposure and following wasp removal. The oviposition depression following wasp exposure is maintained for an extended period following wasp removal. Of unique interest is that this behavior is dependent on learning and memory genes. This genetic identification demonstrates that the behavior is indeed a *bona fide* long-term memory paradigm(Kacsoh *et al.* 2018, e1007430; Kacsoh *et al.* 2015c, 10.7554/eLife.07423; Kacsoh *et al.* 2017; Lefevre *et al.* 2012, 230-233; Lynch *et al.* 2016). We performed an egg depression assay similar to previously published protocols (see methods). Briefly, flies were CO_2_ anesthetized and sorted by genotype to ensure that both the GAL4 and hairpin are in the same genetic background. Flies were then sorted into groups of 10 females and 5 males. Unexposed flies are placed into standard Drosophila vials containing 5 mL standard Drosophila media. For exposed groups, batches of 10 female, 5 male flies are placed into standard Drosophila vials with 20 female wasps. Following a 24-hour exposure period (acute response), flies from both unexposed and exposed conditions are briefly, but independently anesthetized. Unexposed flies were placed into new vials. Exposed flies were separated from wasps following anesthetization. These wasp-exposed flies were placed into new vials. Eggs were counted from the vials that flies were removed from, which corresponds to the acute response, 0-24-hour period. After an additional 24-hours in new vials, all flies were removed from all treatment vials and eggs were counted to measure the memory response (24-48-hour period) (Figure 2 A). We performed all experiments at 25°C with a 12:12 light:dark cycle at light intensity 16_7_, using twelve replicates at 40% humidity. To avoid bias, prediction scores for the genes being tested remained unknown until the completion of the study. A random number was assigned to each stock and the only genotypic information provided was the presence/absence of a balancer chromosome. All vials were coded and scoring was blind as the individual counting eggs was not aware of treatments or genotypes. Eggs were counted immediately after fly removal for both days tested to measure acute and memory responses.

**Figure 2.**
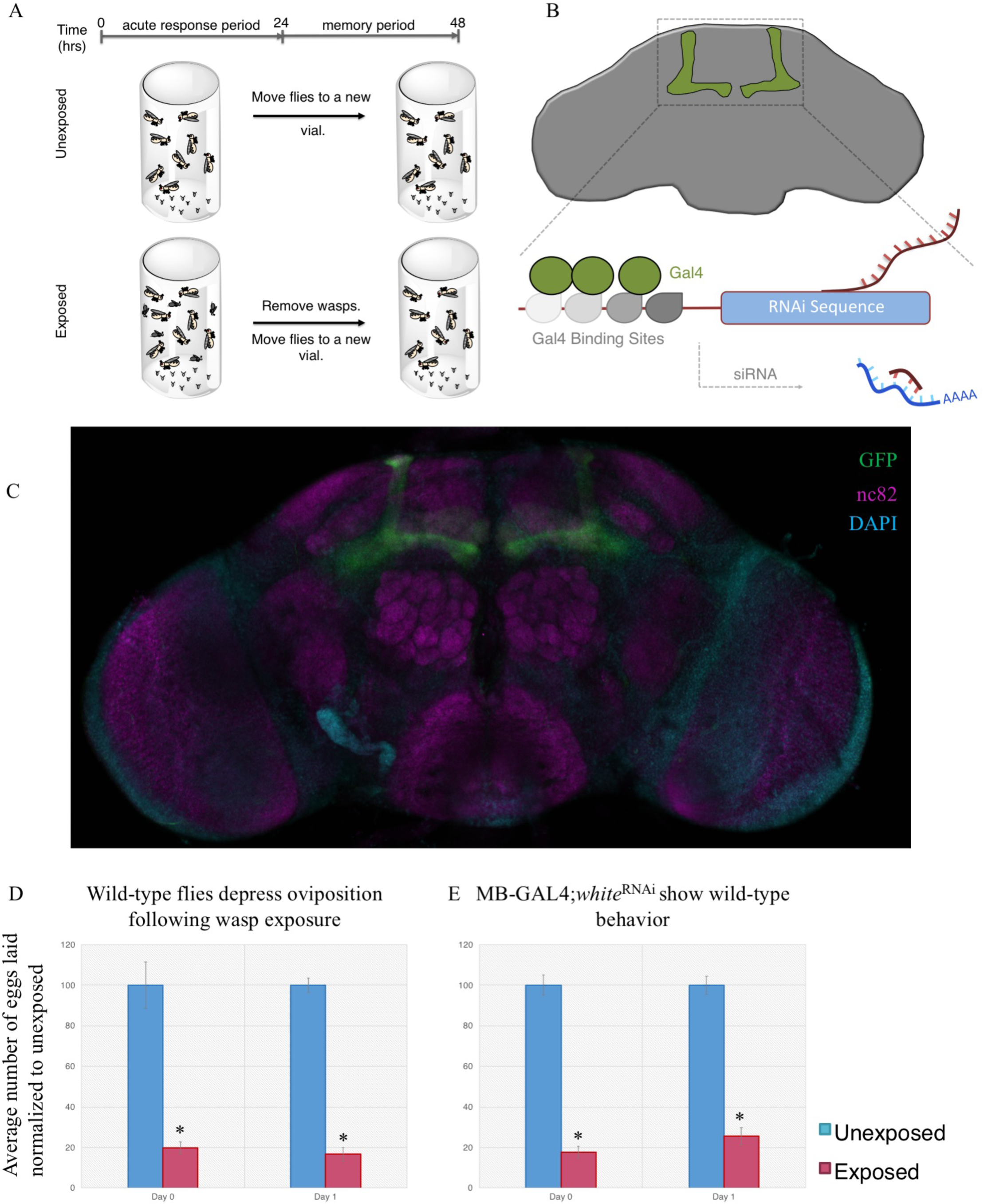
Wild-type flies depress oviposition following wasp exposure. Experimental design for oviposition depression assay (A) indicates a 0-24-hour period while wasps are present (acute exposure) and a 24-48-hour period following wasp removal (memory). In order to screen for long term memory genes, we drive expression of various RNAi hairpin constructs in the fly mushroom body (MB) (B). A merged image of the MB247 driver line expressing a CD8-GFP in green, DAPI in teal, and nc82 in magenta is shown in (C). Using this assay, we present the percentage of eggs laid by exposed flies normalized to eggs laid by unexposed flies. Wild-type flies exposed to wasps lay fewer eggs than unexposed flies in the presence of and following wasp removal (D). Flies expressing a white^RNAi^in the MB also show wild-type behavior, indicating that the RNAi machinery does not perturb the acute response nor the memory response (E). Error bars represent standard error (n = 12 biological replicates) (*p < 0.05).

We first tested wild-type flies, followed by utilization the yeast based GAL4/UAS system. The GAL4/UAS system allows us to drive the expression of a genetic construct, in this case RNA-hairpins targeting each of the predicted gene mRNAs. These RNAi-hairpins were in conjunction with the MB driver MB247 (Supplementary Table 1) (Figure 2 B) (Roman *et al.* 2001, 12602-12607; Duffy 2002, 1-15). The MB247 driver allows us to perturb mRNA levels in a given tissue only (MB) and not the entire organism (Figure 2 C, Supplementary Figure 2). This specificity allows us to delineate the role of the gene in the learning and memory center of the fly brain from the rest of its roles in the organism. As previously shown, we find that wild-type flies depress oviposition during the acute response period (in the presence of wasps) and maintain the depression during the memory response period (following wasp removal) (Figure 2 D) (Lefevre *et al.* 2012, 230-233; Lynch *et al.* 2016; Kacsoh *et al.* 2018, e1007430; Kacsoh *et al.* 2015c, 10.7554/eLife.07423; Kacsoh *et al.* 2017). We also reconfirm that the driver line alone (outcrossed to *Canton* S) has wild-type memory retention (Supplementary Figure 1) (Kacsoh *et al.* 2015c, 10.7554/eLife.07423; Kacsoh *et al.* 2017). We tested an additional control line, where we expressed an RNA-hairpin targeting the white gene in the MB. This genetic combination yielded wild-type behavior, which demonstrates that induction of a RNA-hairpin construct alone does not induce a deficient acute response or memory formation (Figure 2 E).

Using this system as a means to probe stimulus response and memory, we utilized the CAFA analysis to provide a series of potential long-term memory genes (Table 1). We then used the GAL4/UAS system to drive expression of an RNA-hairpin targeting each of the predicted gene mRNAs in conjunction with the MB driver. Our assay allows us to measure the role of each of these genes for the acute response and memory response.

**Table 1.**
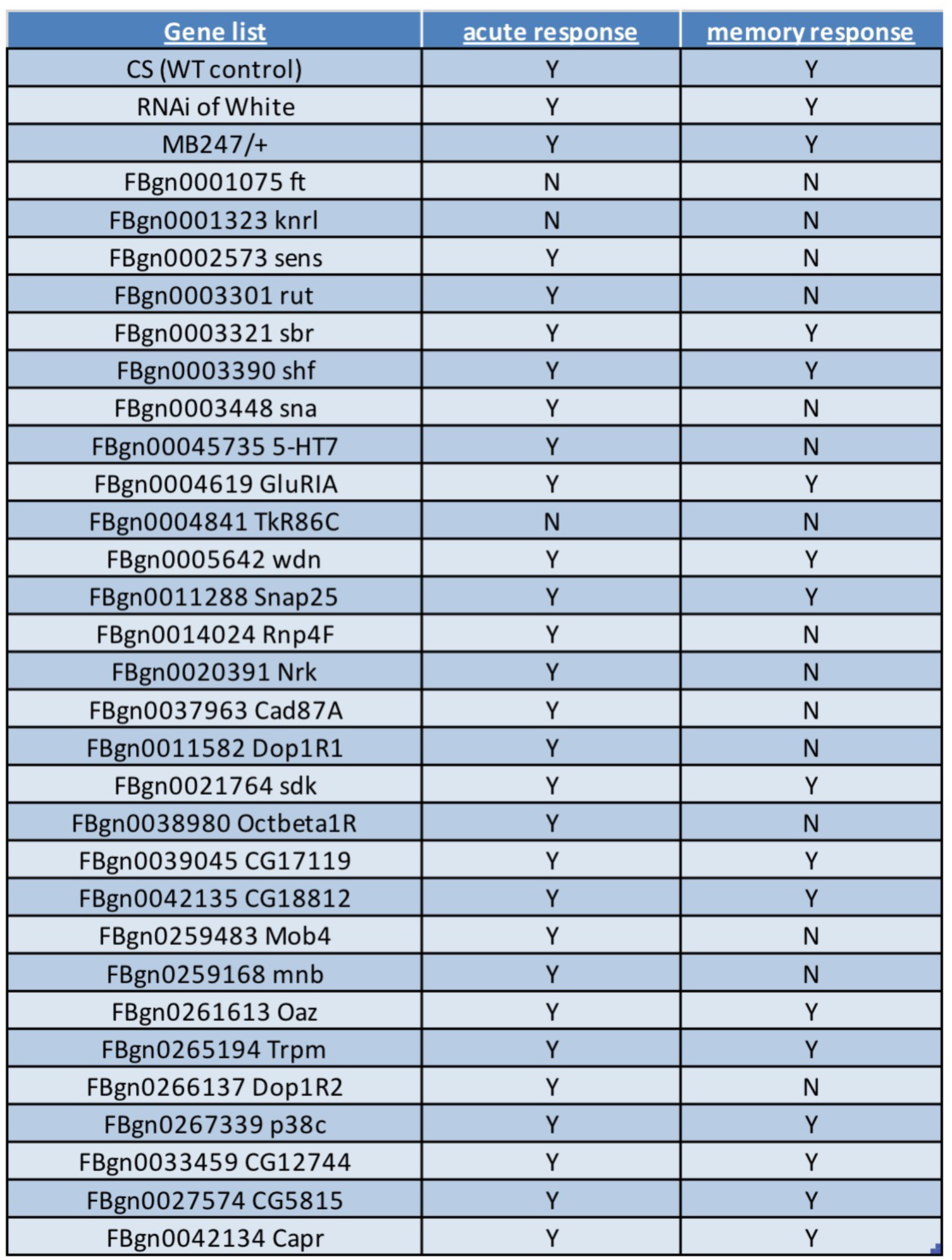
Bioinformatic analysis reveals a series of genes that may be involved in long-term memory formation. Genes tested and results are shown. A binary Yes (Y) / No (N) output is shown for each gene with respect to the presence of an acute response and a memory response. Both an acute and memory phenotype are listed if the egg count is not significantly different from unexposed. Genes that perturb the acute response might be involved in memory formation, but their function in memory cannot be teased apart in the manner tested.

### THREE CAFA-SELECTED GENES AFFECT IN THE ACUTE RESPONSE

We tested a total of 29 unique RNAi hairpins against 29 unique mRNA species in the MB and found 3 genes that, when knocked down via the expression of an RNAi transgene, elicit no acute response. These genes are fat (*ft*), knirps-like (*knrl*), and tachykinin-like receptor at 86C (TkR86C) (Figure 3). Fat is a tumor suppressor gene that encodes a large cadherine transmembrane protein. Additionally, it functions as a receptor in the Hippo signaling pathway and the Dachsous-Fat planar cell polarity pathway(Matakatsu and Blair 2006, 2315-2324; Matakatsu and Blair 2004, 3785-3794). It has GO annotation enrichments in cell cycle, development, and response to stimuli. Interestingly, *ft* has been implicated in neurodegenerative disorders that are polyglutamine (ployQ) diseases, where a functional Hippo/Fat pathway is protective(Napoletano *et al.* 2011, 945-958). The acute response may have been perturbed due to early onset neurodegeneration as a result of the *ft* knockdown (Figure 3 A).

**Figure 3.**
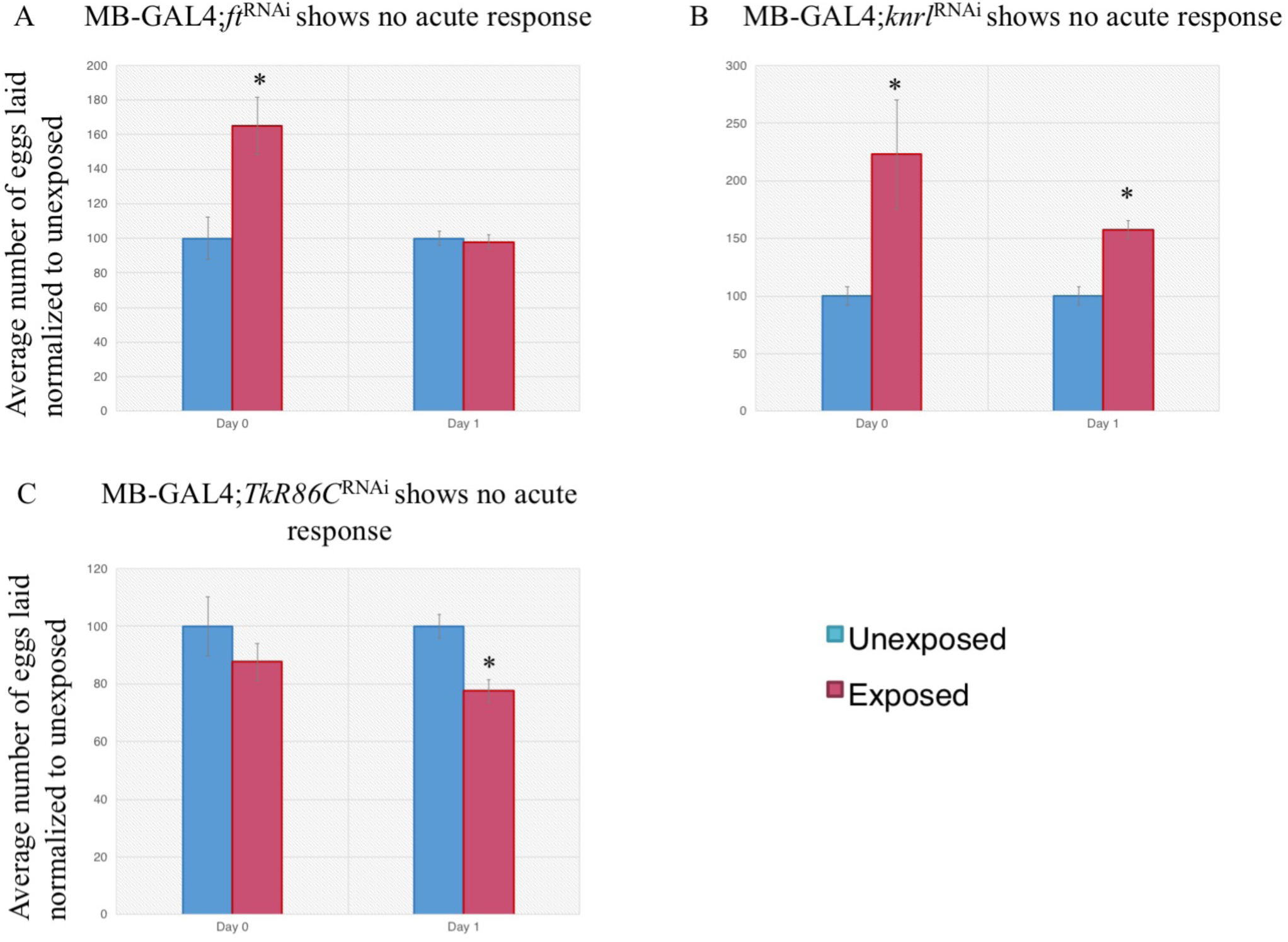
Expression of several genes in the mushroom body is necessary for the acute response. Percentage of eggs laid by exposed flies normalized to eggs laid by unexposed flies is shown. Flies exposed to wasps with various RNAi hairpins expressed in the MB show no acute response for ft^RNAi^(A), knrl^RNAi^(B), and TkR86C^RNAi^(C). Error bars represent standard error (n = 12 biological replicates) (*p < 0.05).

Knirps-like is a nuclear hormone receptor with a C4 zinc finger motif, lacking a ligand-binding domain. It is a target of hedgehog, wingless, and the Notch signaling pathways, with strong GO term enrichment in development(Chen *et al.* 1998, 4959-4968; Chen 1999; Curators 2004)(Figure 3 B).

TKR86C is a G-protein coupled receptor and is rhodopsin-like. Its molecular function is described as neuropeptide receptor activity and is involved in many biological processes such as neuropeptide signaling and inter-male aggression(Jiang *et al.* 2013, E3526-34; Johnson *et al.* 2003, 52172-52178; Asahina *et al.* 2014, 221-235; Poels *et al.* 2009, 545-556). The data suggest that TKR86C may be involved in the acute response, perhaps by way of neuropeptide signaling (Figure 3 C, Table 1).

### TWELVE CAFA-SELECTED GENES AFFECT IN THE MEMORY RESPONSE

Of the total 29 RNAi hairpins against unique mRNA species in the MB tested, we identified a total of 12 unique mRNA species that are required in the MB for long-term memory formation: senseless (*sens*), rutabaga (*rut*), snail (*sna*), 5-hydroxytrptamine (serotonin) receptor 7 (*5-HT7*), RNA-binding protein 4F (*Rnp4F*), Neuro-specific receptor kinase (*Nrk*), Cadherin 87A (*Cad87A*), Dopamine 1-like receptor 1 (*Dop1R1*), Octopamine β1 receptor (*Octβ1R*), MOB kinase activator 4 (*Mob4*), Minibrain (*mnb*), and Dopamine-like receptor 2 (*Dop1R2*) (Figure 4). Previous work utilizing this assay has implicated the MB in long-term memory formation (Kacsoh *et al.* 2015c, 10.7554/eLife.07423), and given our results in this study, we have now identified additional gene products that also modulate the formation of this long-term memory.

**Figure 4.**
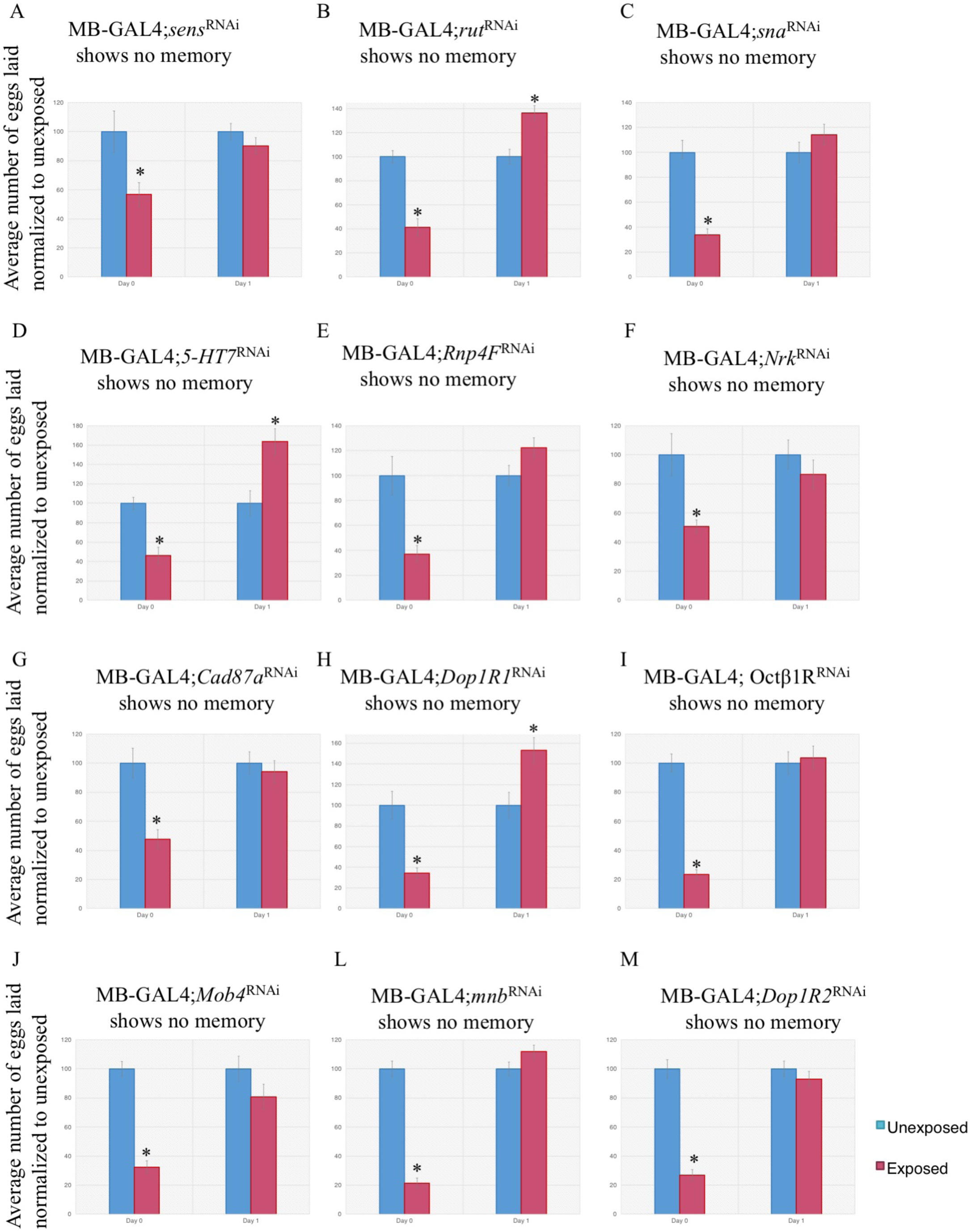
Expression of several genes in the mushroom body is necessary for long-term memory formation. Percentage of eggs laid by exposed flies normalized to eggs laid by unexposed flies is shown. Flies exposed to wasps with various RNAi hairpins expressed in the MB show no memory response for sens^RNAi^(A), rut^RNAi^(B), sna^RNAi^(C), 5-HT7^RNAi^(D), Rnp4F^RNAi^(E), Nrk^RNAi^(F), Cad87A^RNAi^(G), Dop1R1^RNAi^(H), Octβ1R^RNAi^(I), Mob4^RNAi^(J), mnb^RNAi^(K), and Dop1R2^RNAi^(L). Error bars represent standard error (n = 12 biological replicates) (*p < 0.05).

We identified 12 genes involved in long-term memory formation, 11 of which are novel members of long-term memory formation using this paradigm. We define a gene to be a candidate for long-term memory formation if there is a statistically significant egg depression during the acute response, but the depression is no longer statistically significant following wasp removal. *Sens* is a transcription factor that stimulates expression of pro-neural genes by regulating the differentiation of the peripheral nervous system and the R8 photoreceptor (Figure 4 A). It is GO term enriched in development and in responses to stimuli(Frankfort and Mardon 2004, 563-570; Salzberg *et al.* 1994, 269-288; Morey *et al.* 2008, 795; Mishra *et al.* 2013, e1004027). *Rut* encodes a membrane bound calcium^2+^/calmodulin activated adenylyl cyclase that is responsible for the synthesis of cAMP(Levin *et al.* 1992, 479-489). It is GO term enriched in behavior, responses to stimuli, and nervous system processes. *Rut* has been implicated in long-term memory formation before in associative learning(McGuire *et al.* 2003, 1765-1768; Zars *et al.* 2000, 18-31; Dubnau *et al.* 2001b, 476-480; Alonso *et al.* 2003, 653-657; Margulies *et al.* 2005, R700-R713; Qin and Dubnau 2010, 203-212), as well as in non-associative learning using the fly-wasp assay(Kacsoh *et al.* 2015a, 1143-1157; Kacsoh *et al.* 2015c, 10.7554/eLife.07423), thus further validating our experimental protocol and results (Figure 4 B). *Sna* is a transcription factor that contributes to embryonic mesoderm development and nervous system development(Ashraf *et al.* 1999, 6426-6438; Mathew *et al.* 2011, 4978-4993). It is GO term enriched in development (Figure 4 C). *5-HT7* is a serotonin G-protein-coupled-receptor that binds and transmits the signal from the neurotransmitter 5-HT (serotonin) (Gaudet *et al.* 2010; Gasque *et al.* 2013, srep02120) and has been shown to be involved in olfactory learning and memory(Becnel *et al.* 2011, e20800; Luo *et al.* 2012, 471-484). It is GO term enriched in behavior and responses to stimuli (Figure 4 D). *Rnp4F* is predicted to be involved in mRNA binding, but little has been done to elucidate its function(Curators 2004; Lasko 2000, F51-6) (Figure 4 E). *Nrk* is a receptor tyrosine kinase that is expressed during neuronal development and regulates axon path finding(Kucherenko *et al.* 2011, 228-242; Oishi *et al.* 1997, 11916-11923; Marrone *et al.* 2011, 1). It is GO term enriched in development and response to stimulus (Figure 4 F). *Cad87A* is currently known to be cadherin-like, but its function and biological processes are unknown(Curators 2004) (Figure 4 G). *Dop1R1* is known to be involved in dopamine neurotransmitter activity and visual learning and memory, not unlike the assay we utilized(Gotzes *et al.* 1994, 131-141; Seugnet *et al.* 2008, 1110-1117; Brody and Cravchik 2000, F83-8). Its GO term enriched in nervous system processes, behavior, responses to stimuli, and signaling. Interestingly, mutations in the human ortholog has been shown to correlate with attention deficit disorder(Lowe *et al.* 2004, 348-356) (Figure 4 H). *Octβ1R* is known to be a G-protein-coupled-receptor and is rhodopsin-like. It has been shown to have a role in the negative regulation of synaptic growth at the neuro-muscular-junction(Balfanz *et al.* 2005, 440-451; Brody and Cravchik 2000, F83-8; Koon and Budnik 2012, 6312-6322). It is GO term enriched in responses to stimuli and signaling (Figure 4 I). *Mob4* is involved in the negative regulation of synaptic growth at neuromuscular junctions, neuron projection morphogenesis, axo-dendritic transport, and negative regulation of the hippo pathway(He *et al.* 2005, 4139-4152; Schulte *et al.* 2010, 5189-5203; Zheng *et al.* 2017, 3612-3623). It is GO term enriched in cell cycle, development, responses to stimuli, and signaling (Figure 4 J). *Mnb* is a serine/threonine protein kinase and interacts with multiple signaling pathways. It is involved in multiple facets of behavior, and mutants have deficits in behaviors including visual learning and memory. The mutant has a morphologically smaller brain when compared to wild-type(Degoutin *et al.* 2013, 1176; Helfrich 1986, 321-343; Shaikh *et al.* 2016, 3195-3205; Tejedor *et al.* 1995, 287-301; Sokolowski 2001, 879). It is GO term enriched in development, nervous system process, behavior,responses to stimuli, and signaling. Human orthologs exit of this gene (DYRK1A) and manifests as mental retardation(Møller *et al.* 2008, 1165-1170) (Figure 4 K). *Dop1R2* belongs to the g-protein-coupled-receptor family and is implicated in adrenergic receptor activity, dopamine neurotransmitter activity, positive regulation of oxidative stress-induced neuron death, and cell signaling. These phenotypes are generally localized to the fanshaped body of the brain, which is the visual learning and memory center of the fly brain that projects to the MB(Gaudet *et al.* 2010; Brody and Cravchik 2000, F83-8; Han *et al.* 1996, 1127-1135; Radford *et al.* 2002, 38810-38817; Cohn *et al.* 2015, 1742-1755; Pimentel *et al.* 2016, 333). This gene is GO term enriched in behavior, responses to stimuli, and signaling (Figure 4 L).

An additional 14 lines were tested and demonstrate wild-type, statistically significant, acute and memory responses during and following wasp exposure (Supplementary Figure 3). These results further demonstrate that knockdown of any gene does not result in a perturbation— knockdown where we observe a perturbation of wild-type behavior does implicate said gene/gene product in the acute or memory process.

### GENE NETWORK ANALYSIS REVEALS A HIGH DEGREE OF CONNECTIVITY

In order to further examine both the acute response genes and the memory genes identified from our screen, we employed the integrative <u>m</u>ulti-species <u>p</u>rediction (IMP) webserver to determine the extent to which our hits are connected with either other response to stimulus genes or to known learning and memory genes(Wong *et al.* 2015, W128-33; Wong *et al.* 2012, W484-90). IMP is a user-friendly program that provides the integration of gene-pathway annotations from a selected organism (in our case, *D. melanogaster*), and subsequently maps functional analogs by providing a userfriendly visual output. IMP has been utilized to predict gene-process predictions that were subsequently experimentally validated(Chikina *et al.* 2009, e1000417; Chikina and Troyanskaya 2011, e1001074; Kacsoh *et al.* 2017).

We first queried the network with our acute response implicated genes. We set the network filter to only show interactions whose minimum relationship confidence was 0.85, which is a stringent threshold (Figure 5 A). We find that *ft* and *knrl* are connected in a large gene network, while *TkR86C* is only connected to one partner, l(1)G0020. The genes we identified have independently associated GO terms that are predicted with a 0.85 confidence score (Figure 5 B). Interestingly, *ft* is associated with R3/R4 cell differentiation, raising the possibility of perturbed vision in these animals accounting for the lack of the acute response. We find that the network is enriched for the biological processes of cell fate specification (34.6%, p-value: 1.93e-8), cell fate commitment involved in pattern formation (19.2%, p-value: 9.21e-5), skin development (19.2%, p-value: 6.33e-4), and neuroblast differentiation (15.4%, p-value: 6.29e-4).

**Figure 5.**
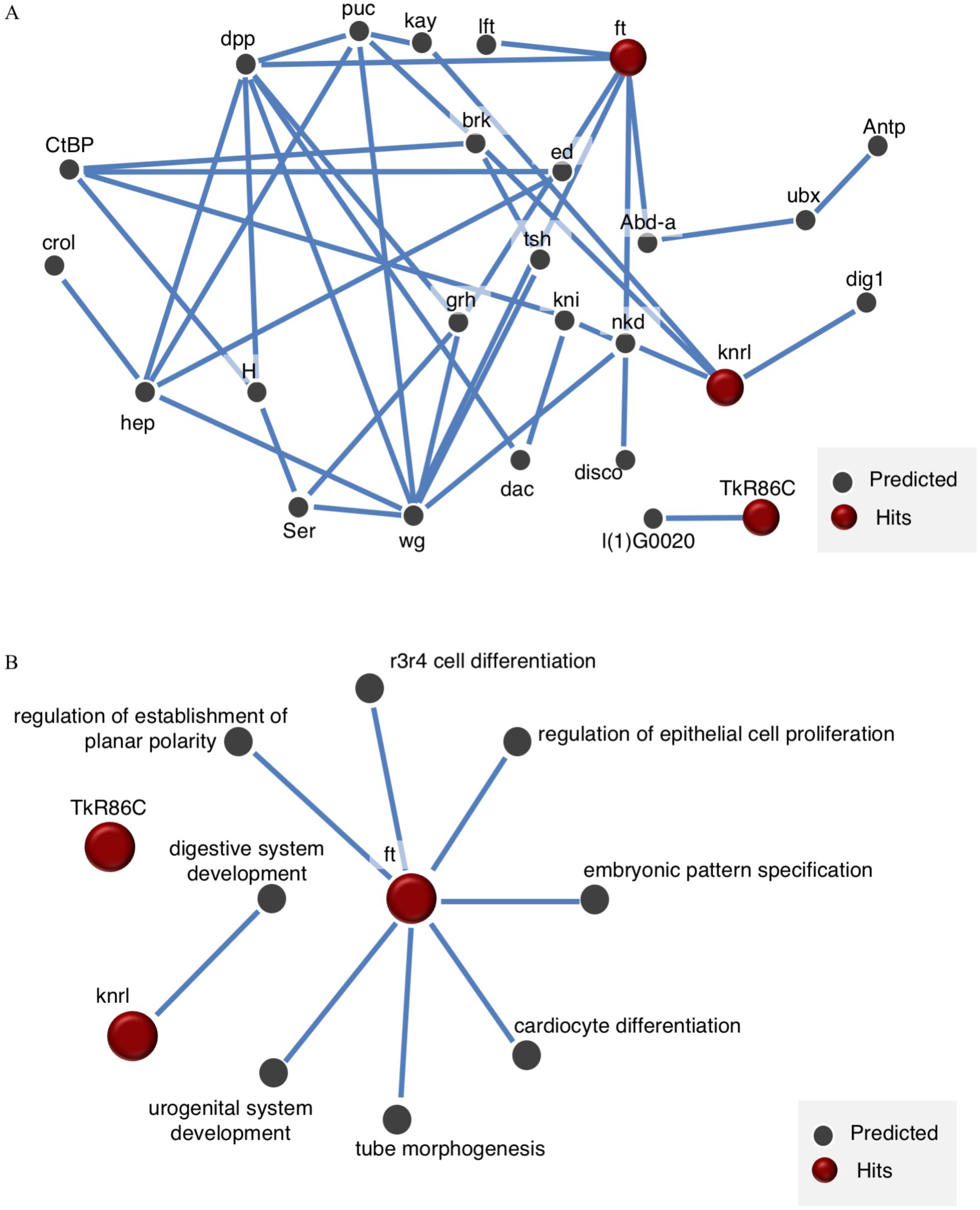
IMP analysis of genes that yielded an acute response defect reveals connectivity and developmental functions. A visual representation of a gene network utilizing our positive acute response hits (ft, knrl, TkR86C) reveals a high level of connectivity between identified genes (A). Lines indicate a minimum relationship confidence of 0.85. GO term enrichment for acute response candidates reveal development specific processes (B).

We next queried the network with our memory response implicated genes. We set the network filter to only show interactions whose minimum relationship confidence was 0.85 (shown in blue), followed by a filter of 0.4 (shown in pink), in order to highlight strong and weak, but relevant, interactions (Figure 6 A). We find that almost every gene tested is highly interconnected. Interestingly, the hits were connected, both directly and indirectly, to known learning and memory genes that have been tested using this paradigm. These genes are FMR1, dunce, amnesiac, alcohol dehydrogenase factor 1, and grunge(Kacsoh *et al.* 2015c, 10.7554/eLife.07423; Kacsoh *et al.* 2015a, 1143-1157; Kacsoh *et al.* 2017; Kacsoh *et al.* 2013, 947-950). Given this level of interconnectivity with previously identified memory genes, the network suggests that we have identified potential *bona fide* learning and memory genes. The GO terms associated with the genes identified are very heavily connected to long-term memory, synaptic functions, and behavior (Figure 6 B). The network is enriched for the biological processes of associative learning (19.4%, p-value: 3.14e-5), learning (19.4%, p-value: 5.69e-5), positive regulation of growth (19.4%, p-value: 6.21e-5), and olfactory learning (16.7%, p-value: 1.27e-4). Only one of the hits, *Nrk*, was found to not be connected to the network. However, a connection was observed with multiple genes following a lowering of the relationship confidence score, indicating that it may have some interaction with the network.

**Figure 6.**
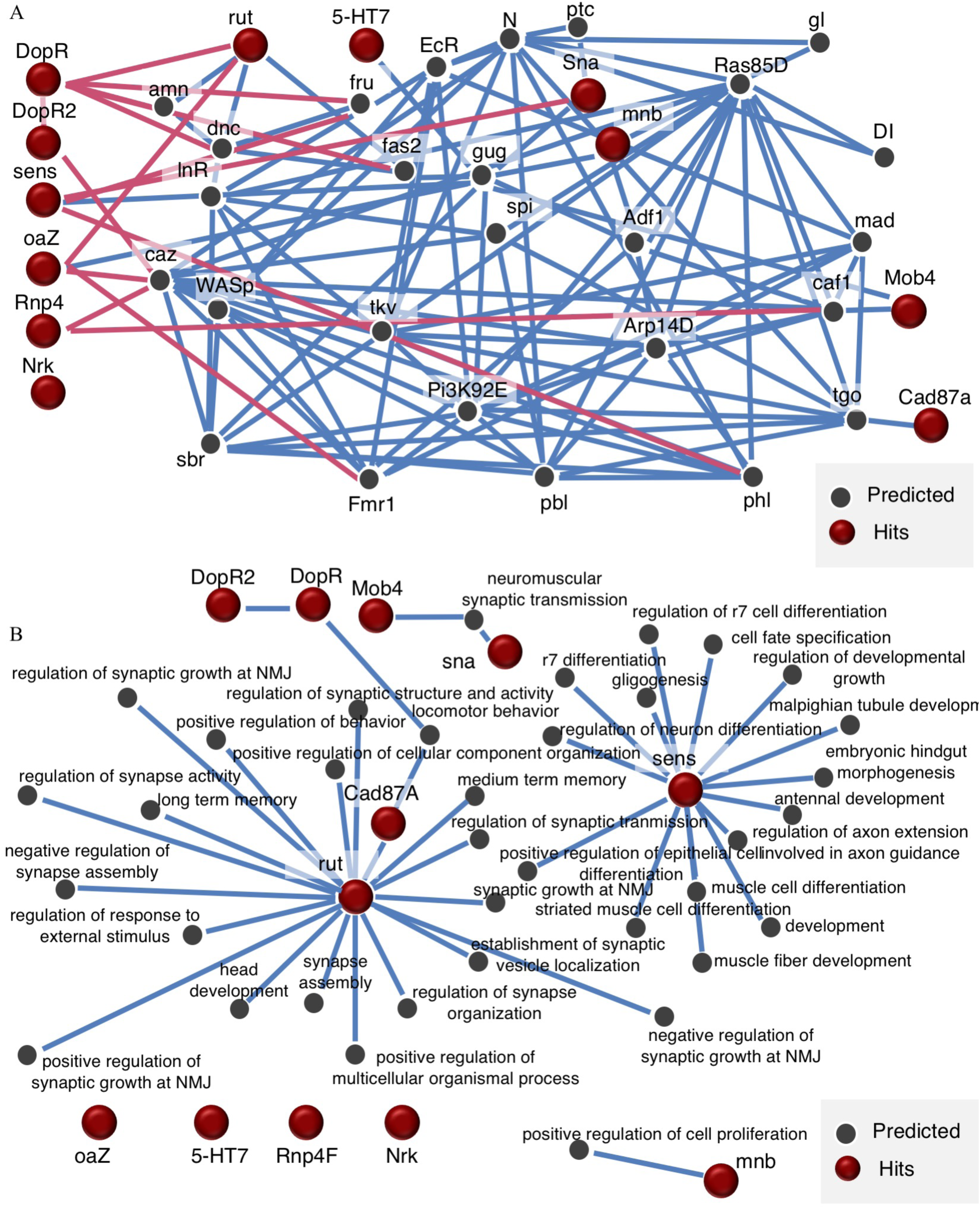
IMP analysis of genes that yielded a memory defect reveals a high level of connectivity in addition to neuro-specific functions. A visual representation of a gene network utilizing our positive long-term memory hits (sens, rut, sna, 5-HT7, Rnp4F, Nrk, Cad87A, Dop161, Octbeta1R, Mob4, mnb, Dop1R2) reveals a high level of connectivity between some of the selected genes (A). Blue lines indicate a minimum relationship confidence of 0.8, while pink lines indicate a relationship confidence of 0.4. GO term enrichment scores for long-term memory candidates reveal neuro specific processes (B).

## DISCUSSION

In this study, we identify novel long-term memory genes as part of the CAFA3 experiment. We analyzed hits for the long-term memory process in Drosophila using a functional network analysis tool, IMP. We identified 3 genes involved in the acute response to wasps, and 12 genes implicated in long-term memory. To our knowledge, our acute response hits and 9 of our genes implicated in long-term memory have not been previously implicated to function in Drosophila memory. Of the remaining 3 genes, 2 have not been shown to be involved in this form of Drosophila memory.

Past work has demonstrated the need for an intact visual system in order to elicit a fly behavioral response in the presence of the wasp (Kacsoh *et al.* 2017; Kacsoh *et al.* 2015a, 1143-1157; Kacsoh *et al.* 2015c, 10.7554/eLife.07423; Kacsoh *et al.* 2013, 947-950; Kacsoh *et al.* 2018, e1007430). In this study, we provided the opportunity of direct interaction between the flies and wasps, permitting multi-sensory information acquisition. Using this setup, we identified 3 genes that are necessary in the MB to elicit the oviposition behavior. Thus, it becomes possible that these genes may affect certain sensory components of the nervous system in addition to altering neuronal excitability of the MB. For example, the excitability threshold in the MB could be raised or lowered, changing the responses observed in our behavioral paradigm, not because the gene is involved in the acute response or memory, but rather because the gene is altering neuronal properties. A wild-type fly might achieve maximal excitation and form a stable memory from the wasp interaction, but flies deficient in the 3 acute response genes identified are trained to a lesser degree resulting in an unstable ability to process the information. Following wasp removal, this instability might manifest as a lack of a memory formed. Future experiments that utilize isolated sensory stimuli, especially visual cues given the involvement of photoreceptor cell determination, would be able to examine these hypotheses.

We highlight that stimulus response is not affected in flies lacking our 12 memory hits, as the day 0 (acute response) data is indistinguishable from wild-type levels. The presence of the acute response in these flies suggests that wasp identifying sensory systems are intact. Thus, RNAi of these genes in the MB gives us F1 animals that can response to wasps as wild-type, but forget the exposure following wasp removal. Therefore, the genes that we define as a long-term memory hit are not required for the detection the input of “wasp”, but instead are required to make the initial memory. These results suggest that the wiring and development of the fly brain during knock-down of these select genes remains intact and produces a functioning brain that can respond to the initial stimulus as the wild-type controls (day 0). We observe phenotypic differences on day 1 (memory period), where we find the rapid extinction of memory, or the lack of memory formation. These two results are not possible to delineate with the current experimental setup, but remains as a future question. Wild-type flies demonstrate a maintained oviposition response, demonstrating memory retention following the wasp exposure (Kacsoh *et al.* 2017; Kacsoh *et al.* 2015c, 10.7554/eLife.07423).

It remains a possibility that the genes presented as hits are not only involved in either the acute response or memory formation, but could be participating in development. The MB driver (MB247) in conjunction with each of the RNAi transgenes that presented either an acute response defect or a memory retention phenotype could be the result of the constitutive expression of the RNAi hairpin throughout development and might have perturbed the development, whether morphological or excitatory element of the neuron, yielding the alterations in behavior. This possibility suggests that knockdown phenotypes have the potential to reveal undetected changes that preclude proper adult MB functions. This possibility does not preclude the potential hits we uncovered, nor the efficacy of the prediction tool, but instead provides an important avenue for future experiments.

Finally, we graded our behavioral responses, the acute and memory responses, on a binary scale (yes/no) determined by significance testing when compared to control. This type of comparison does not account for subtle changes that could be contributed by these genes, such as an attenuated initial or learned response, or a heightened initial or learned response. To elucidate these different alternatives, additional testing will be required, utilizing multiple RNAi transgenes targeting a single gene and examining the gene doses.

Given our data, we propose a model where the initial wasp stimulus is partially mediated by genetic factors that could mediate development and/or excitability of the wasp response circuit. These factors, including other yet to be discovered players, to mediate the oviposition depression, which results in the activation of effector caspases(Kacsoh *et al.* 2018, e1007430; Kacsoh *et al.* 2015c, 10.7554/eLife.07423). The maintenance of this physiological change, and formation of a long-term memory, is mediated by a second group of genes that perpetuate the signal so that the flies continue to lay fewer eggs even after removal of the wasp threat (Figure 7).

**Figure 7.**
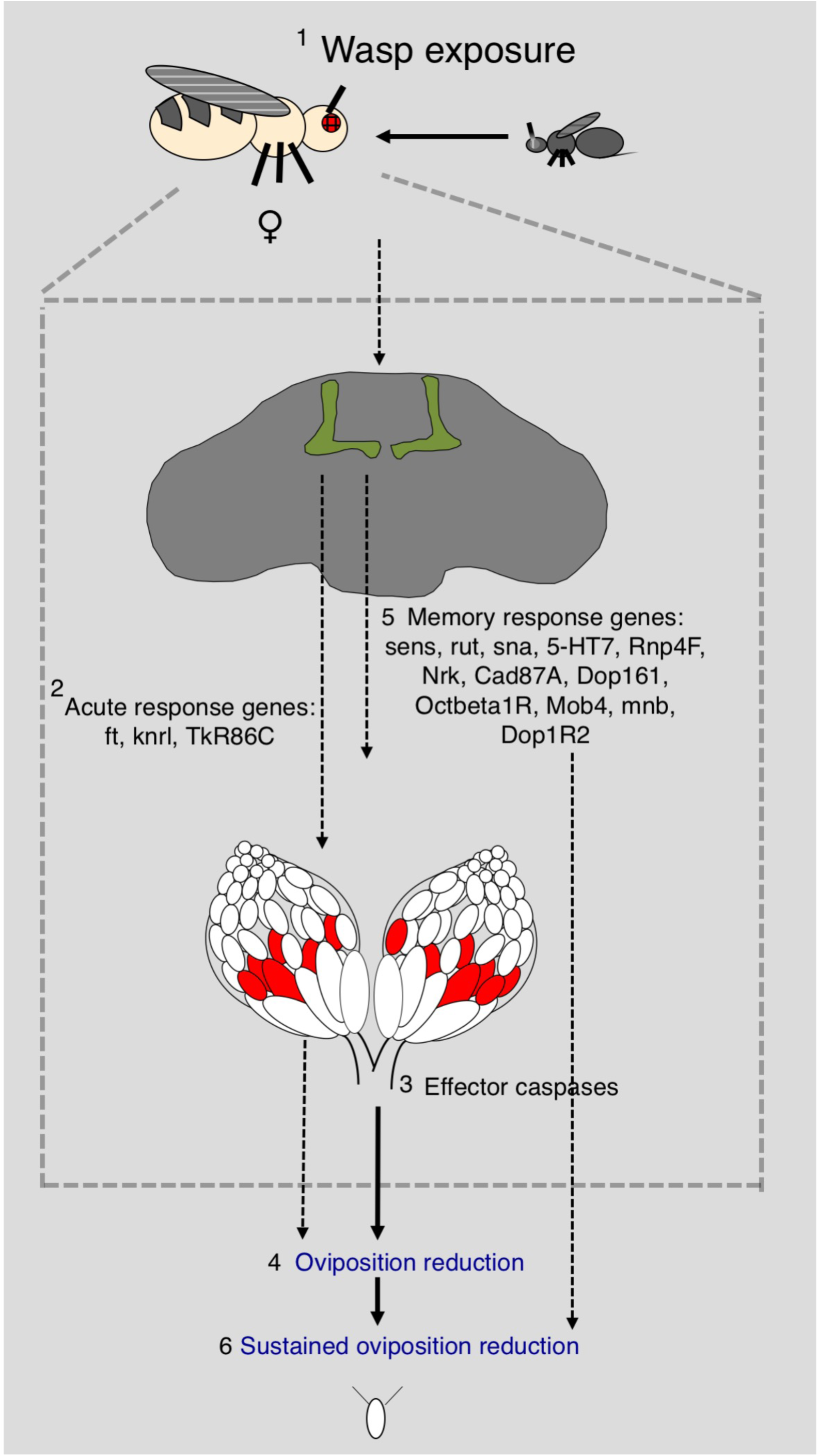
Proposed model for genes identified in the acute and memory response. Model for acute response and learning/memory. First, a female fly sees a wasp. Second, acute response genes are either needed during development of the MB or to initiate a wasp response. Third, effector caspases are activated. Fourth, oviposition depression is observed. Fifth, we see activation of memory genes. Sixth, we see a sustained reduction in oviposition. We hypothesize that some genes are involved in either development of the MB and/or processing in the MB during the acute response, while other genes are activated later and are involved in long-term memory formation. Following memory consolidation, these genes may be further expressed while the initial acute response genes might be dispensable.

Collectively, our data highlights the importance of incorporating bioinformatics-based approaches as a means to identify new genetic factors in phenotypes of interest and at the same time evaluate prediction tools. Some of the genes that we identified have human disease orthologs and thus, we are provided with the opportunity to study these human diseases in a genetic model system, which we are able to show has a phenotype of interest. Gene function annotations, which form the primary means of analysis in CAFA experiments, tend to be biased towards genes with large effect(Hibbs *et al.* 2009, e1000322; Hess *et al.* 2009, e1000407). Pairing these evaluations with targeted experimental assays might elucidate novel regulators and pathways that would have gone otherwise undetected due to detection threshold. These novel elements could have biologically important implications even though they only undergo smaller alterations as a function of a stimulus (Zovkic *et al.* 2013, 61-74). Such changes can be elusive to identify otherwise, yet, could be involved in important biological processes, such as neuron plasticity and excitation. Our study provides a novel role for multiple genes, that have connections to known and novel pathways, as a potential regulator of either a stimulus response, or the maintenance of a response, at the intersection of memory acquisition and memory retention.

## Acknowledgements

We thank Mani Ramaswami, FlyBase, the Vienna Drosophila Resource Center, and the Bloomington Drosophila Stock Center, for stocks. We thank the Developmental Studies Hybridoma Bank at the University of Iowa for reagents. We acknowledge grants from Geisel School of Medicine at Dartmouth (GB and CSG), a Gordon and Betty Moore Foundation Data-Driven Discovery Investigator grant GBMF4552 (CSG), the National Science Foundation grant DBI-1458477, ABI 1458390 (CSG and GB), ABI 1458477 (PR), ABI 1458443 (SDM), and ABI 1458359 (IF), the National Institute of Health Pioneer grant 1DP1MH110234 (GB), and the Defense Advanced Research Projects Agency grant HR0011-15-1-0002 (GB).

## Materials and Methods

### Insect Species/Strains

The *D. melanogaster* strain Canton-S (CS) was used as the wild-type strain, in addition to outcrosses. For crosses:

ft ^RNAi^, knrl ^RNAi^, sens ^RNAi^, rut ^RNAi^, sbr ^RNAi^, shf ^RNAi^, sna ^RNAi^, 5-HT7 ^RNAi^, GluRIA ^RNAi^, TkR86C ^RNAi^, wdn RNAi, Snap25 ^RNAi^, Dop161 ^RNAi^, Rnp4F ^RNAi^, Nrk ^RNAi^, sdk ^RNAi^, CG5815 ^RNAi^, Cad87A ^RNAi^, Octbeta1R ^RNAi^, CG17119 ^RNAi^, Capr ^RNAi^, CG18812 ^RNAi^, mnb ^RNAi^, Mob4 ^RNAi^, Oaz ^RNAi^, Trpm ^RNAi^, Dop1R2 ^RNAi^, p38c ^RNAi^, CG12744RNAi, CG5815 ^RNAi^, and Capr ^RNAi^ were used.

These stocks were acquired from the Bloomington Drosophila Stock Center and the Vienna Drosophila Resource Center (Supplementary Table 1). Stock numbers respectively:

29566, 36664, 27287, 27035, 28924, 55867, 28679, 32471, 27521, 31884, 35654, 27306, 31765, 35457, 55184, 33412, 22203, 28716, 58179, 40823, 110272, 42777, 28888, 65236, 35715, 44503, 51423, 64846 and 67262 from Blooming Drosophila Stock Center and 110272 and 22203 from the Vienna Drosophila Resource Center (Supplementary table 1).

Flies that were aged 4-6 days post-eclosion, grown on fresh Drosophila media, were utilized in all experiments. Flies were maintained at room temperature, approximately 20°C, with a 12:12 light:dark cycle at light intensity 16_7_ with 30-55% humidity dependent on weather. We measured light intensity with a Sekonic L-308DC light meter using shutter speed 120, sensitivity at iso8000, and a 1/10 step measurement value (f-stop). These are conditions used in previous studies(Kacsoh *et al.* 2018, e1007430; Kacsoh *et al.* 2017; Kacsoh *et al.* 2015c, 10.7554/eLife.07423; Kacsoh *et al.* 2015a, 1143-1157).

We utilized the generalist Figitid larval endoparasitoid *Leptopilina heterotoma* (strain Lh14), that is known to infect a wide array of Drosophilids(Schlenke *et al.* 2007, e158; Kacsoh and Schlenke 2012, e34721; Kacsoh *et al.* 2014, 697-715) for all experiments (exposed condition). *L. heterotoma* strain Lh14 originated from a single female collected in Winters, California in 2002, and has been maintained in the lab ever since. To propagate wasp stocks, we used adult *D. melanogaster* (strain *Canton S*) in batches of 40 females and 15 males per vial (Genesse catalog number 32-116). Adult flies are permitted to lay eggs in standard Drosophila vials containing 5 mL standard Drosophila media supplemented with live yeast (approximately 25 granules) for 3-5 days before being removed. Flies are replaced by adult wasps, specifically 15 female and 6 male wasps, for infections. Larval endoparasitoid wasps deposit eggs in developing fly larvae, specifically to the L2 stage of larvae. Wasp containing vials are supplemented with approximately 500 μL of a 50% honey/water solution applied to the inside of the cotton vial plugs. We utilized organic honey as a supplement in all cases. Wasps aged 7 days post eclosion were used for all infections and experiments. During the aging process, female and male wasps were cohabitated, allowing for mating such that no virgin wasps were used. Wasps were never reused for experiments. If wasps were used for an experiment, they were subsequently disposed of and not used to propagate the stock.

### Cross set-up

For RNAi experiments, male flies containing the hairpins were crossed to virgin female flies containing the MB247 GAL4, which were kindly provided by Mani Ramaswami (University of Dublin, Ireland). 10 female MB247 GAL4 virgin flies were mated to 5 males containing the selected hairpin. Crosses were performed in standard Drosophila vials containing 5 mL standard Drosophila media (Genesse catalog number 32-116) supplemented with 10 granules of activated yeast. Crosses were kept at approximately 20°C, with a 12:12 light:dark cycle at light intensity 16_7_ with 30-55% humidity dependent on weather. Crosses were moved to new vials every two days until egg production declined, at which point the adults were disposed of. Upon F1 eclosion, F1 flies were moved to a new vial and allowed to age to 4 days before use in experiments.

### Wasp response and memory assay

F1 flies from above described crosses are CO_2_ anesthetized and sorted by genotype to ensure that both the GAL4 and hairpin are in the same background. Flies are sorted into groups of 10 females and 5 males. Unexposed flies are placed into standard Drosophila vials (Genesse catalog number 32-116) containing 5 mL standard Drosophila media. Once unexposed vials are set up, wasps are anesthetized on the CO_2_ pad next to flies that will be used for exposed vials. Batches of 10 female, 5 male flies are placed into standard Drosophila vials (Genesse catalog number 32-116) with 20 female Lh14 wasps. Vials contain 5mL standard Drosophila media. Following set up, vials are placed into a walk-in incubator with exposed and unexposed vials separated by distance and by vision to prevent social information exchange. Following a 24-hour exposure period (acute response), flies from both unexposed and exposed conditions were briefly, but independently anesthetized. Unexposed flies were placed into new vials containing 5mL standard Drosophila media. Exposed flies were separated from wasps following anesthetization. These wasp-exposed flies were placed into new vials containing 5mL standard Drosophila media. Eggs were counted from the vials that flies were removed from, which corresponds to the acute response, 0-24-hour period. After an additional 24-hours in new vials, all flies were removed from all treatment vials and eggs were counted to measure the memory response (24-48-hour period).

All experimental treatments were run at 25°C with a 12:12 light:dark cycle at light intensity 16_7_, using twelve replicates at 40% humidity unless otherwise noted. To avoid bias, gene categorization following CAFA analysis was withheld until completion of all experiments. Additionally, all vials were coded and scoring was blind as the individual counting eggs was not aware of treatments or genotypes and random subsets were counted by two individuals to ensure accuracy. Eggs were counted immediately after fly removal for both days tested. All raw egg counts and corresponding p-values are provided in supplementary file 1.

### Immunofluorescence

Brains expressing MB247 GAL4 and CD8-GFP were dissected in a manner previously described(Kacsoh *et al.* 2015b, 1143-1157). Flies were placed in batches into standard vials (Genesee catalog number 32-116) of 20 females, 2 males. Three vials were prepared to produce three replicates to account for batch effects. We observed no batch effects. Brains that were prepared for immunofluorescence were fixed in 4% methanol-free formaldehyde in PBS with 0.001% Triton-X for approximately five minutes. The samples were then washed in PBS with 0.1% Triton-X. Samples were then blocked in a 4% NGS solution, prepared in a 0.1% Triton X solution. Brains were stained with nc82 (Hybridoma Bank at the University of Iowa, Antibody registry ID: AB 2314866) at a 1:10 ratio of antibody:0.1% Triton X. Staining was performed overnight at 4C. Following the overnight stain, brains were washed with a 0.1% Triton X solution three times. This was followed by a 10-minute nuclear stain with 4’, 6-diamidino-2-phenylindole (DAPI). Samples were then washed again a 0.1% Triton X solution three times and mounted in Vecta Shield (Vector Laboratories, Catalog number: H-1200).

### Imaging

A Nikon A1R SI Confocal microscope was used for imaging GFP in the MB (Fig 2 C, Supplementary Figure 2). Image averaging of 16x during image capture was used for all images.

### IMP analysis

Following all knockdown experiments, we input our acute response gene hits and memory response gene hits separately into IMP (http://imp.princeton.edu) (Figure 5,6) (Wong *et al.* 2015, W128-33; Wong *et al.* 2012, W484-90). We did this analysis after all experiments were completed to further remove any bias. We utilized a stringent minimum gene connection of 0.85 confidence, indicated with blue lines, as well as a less stringent cutoff of 0.4, indicated with pink lines.

### Statistical analysis

We performed statistical tests on exposed v unexposed conditions in Microsoft Excel. Our conditions utilized Welch’s two-tailed t-tests for all comparisons. P-values reported are included in supplemental file 1.

## SUPPLEMENTARY FIGURE LEGENDS

**Supplementary Figure 1.**
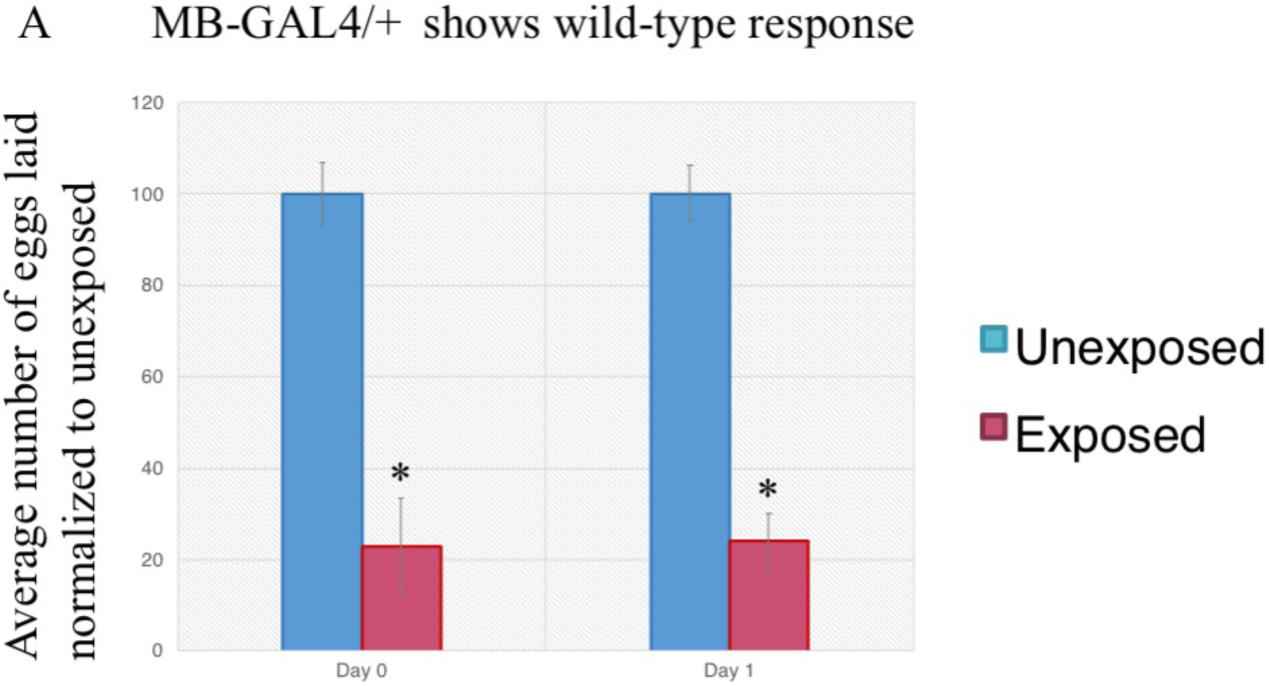
Wild-type flies lay fewer eggs in the presence of wasps. Percentage of eggs laid by exposed flies normalized to eggs laid by unexposed flies is shown. Wild-type flies exposed to wasps lay fewer eggs than unexposed flies for the outcrossed driver line, MB247/+ (A). Error bars represent standard error (n = 12 biological replicates) (*p < 0.05).

**Supplementary Figure 2.**
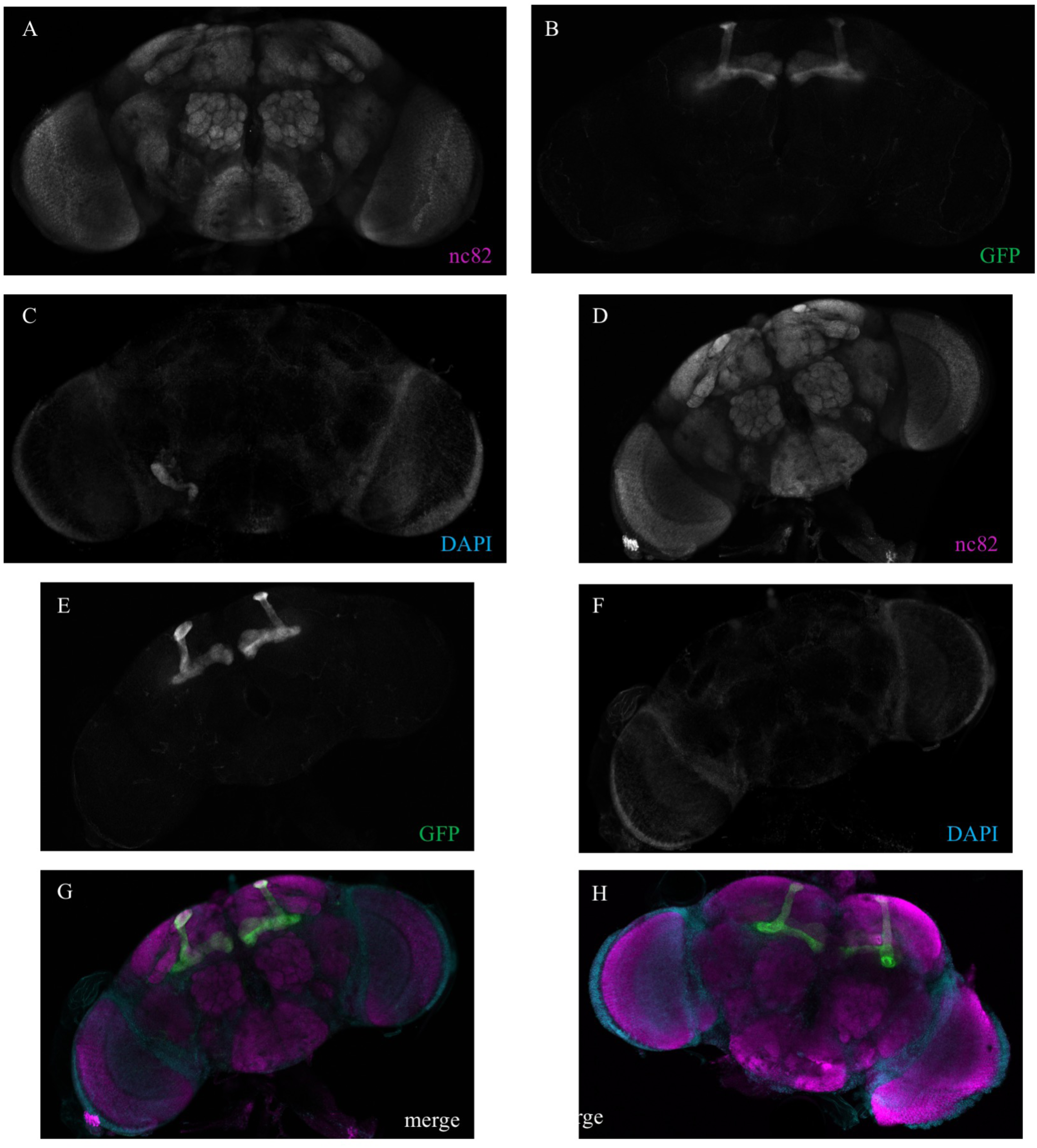
Expression of MB247 driver visualized by CD8-GFP expression. Brains from flies expressing CD8-GFP in conjunction with the MB247 driver line are shown. (A-C) correspond to individual channels of merged image shown in Figure 2 C. Additional brain sample with individual channels and merged image shown in (D-G). An additional brain with only the merged channels is shown in (H). For merged images, teal corresponds to DAPI, magenta corresponds to nc82, and green corresponds to CD8-GFP.

**Supplementary Figure 3.**
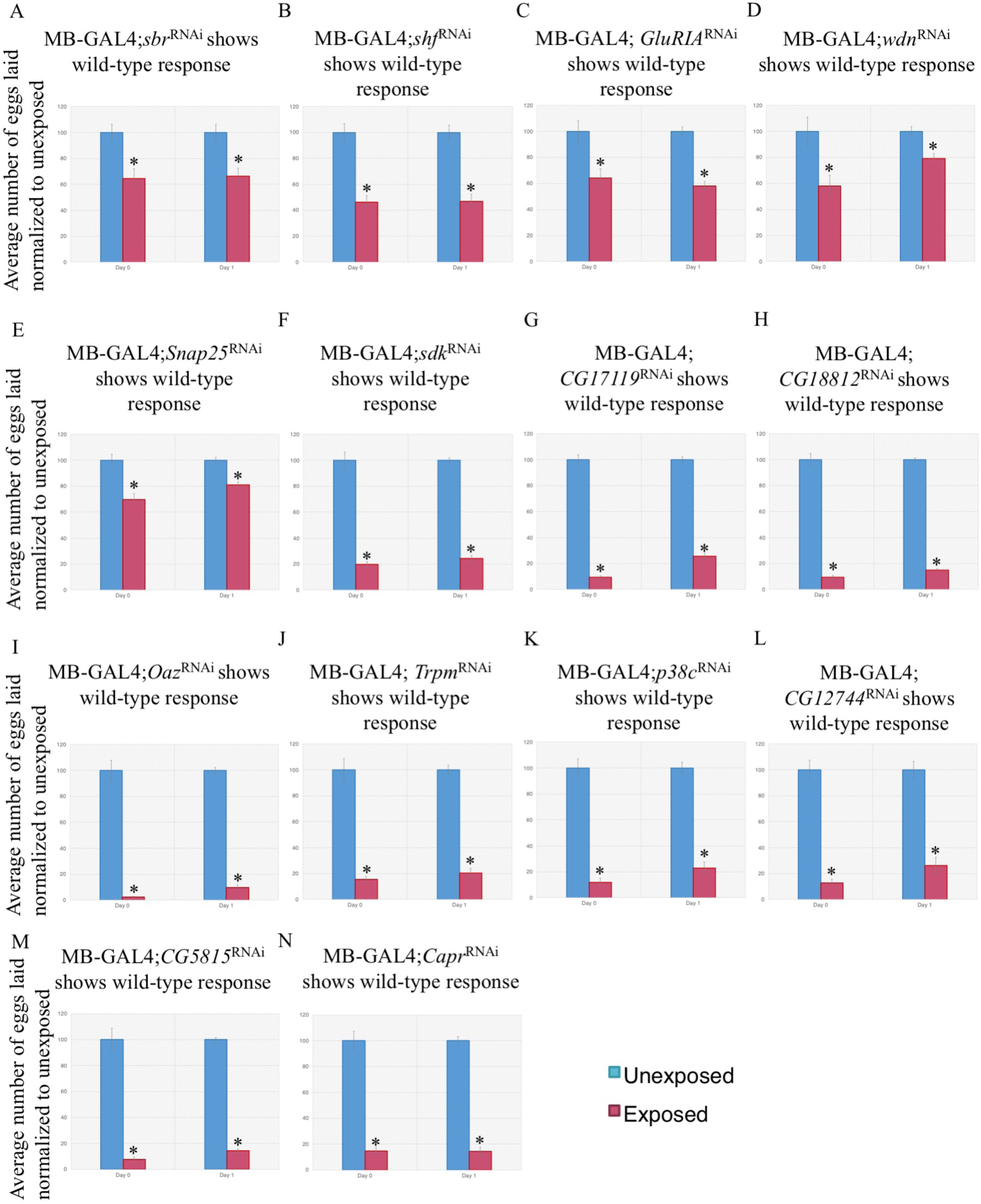
RNAi of genes in the MB show wild-type acute and memory responses. Percentage of eggs laid by exposed flies normalized to eggs laid by unexposed flies is shown. Flies exposed to wasps with various RNAi hairpins expressed in the MB lay fewer eggs than unexposed flies during the acute and memory responses for sbr^RNAi^(A), shf^RNAi^(B), GluRI^RNAi^(C), wdn^RNAi^(D), Snap25^RNAi^(E), sdk^RNAi^(F), CG17119^RNAi^(G), CG18812^RNAi^(H), Oaz^RNAi^(I), Trpm^RNAi^(J),p38c^RNAi^(K), CG12744^RNAi^(L), CG5815^RNAi^(M), and Capr^RNAi^(N). Error bars represent standard error (n = 12 biological replicates) (*p < 0.05).

## Supplementary Table Legends

**Table S1.** Fly genotypes used in this study. Name, genotype, acquisition location, and stock identification number (if applicable) are shown.

| <u>Gene/Allele</u> | <u>Genotype</u> | <u>Acquisition location</u> | <u>ID #</u> |
| --- | --- | --- | --- |
| CS (WT control) | - | n/a | n/a |
| RNAi of White | w[1118]; P{y[+t7.7] w[+mC]=UAS-QS.P}attP2 | Bloomington | 30033 |
| MB247/+ | P{MB247} | Mani Ramaswami | n/a |
| FBgn0001075 ft | y[1] v[1]; P{y[+t7.7] v[+t1.8]=TRiP.JF03245}attP2 | Bloomington | 29566 |
| FBgn0001323 knrl | y[1] sc[*] v[1]; P{y[+t7.7] v[+t1.8]=TRiP.HMS01552}attP2 | Bloomington | 36664 |
| FBgn0002573 sens | y[1] v[1]; P{y[+t7.7] v[+t1.8]=TRiP.JF02575}attP2 | Bloomington | 27287 |
| FBgn0003301 rut | y[1] v[1]; P{y[+t7.7] v[+t1.8]=TRiP.JF02361}attP2/TM3, Sb[1] | Bloomington | 27035 |
| FBgn0003321 sbr | y[1] v[1]; P{y[+t7.7] v[+t1.8]=TRiP.HM05135}attP2 | Bloomington | 28924 |
| FBgn0003390 shf | y[1] sc[*] v[1]; P{y[+t7.7] v[+t1.8]=TRiP.HMC04137}attP2 | Bloomington | 55867 |
| FBgn0003448 sna | y[1] v[1]; P{y[+t7.7] v[+t1.8]=TRiP.JF03094}attP2 | Bloomington | 28679 |
| FBgn00045735 5-HT7 | y[1] sc[*] v[1]; P{y[+t7.7] v[+t1.8]=TRiP.HMS00471}attP2 | Bloomington | 32471 |
| FBgn0004619 GluRIA | y[1] v[1]; P{y[+t7.7] v[+t1.8]=TRiP.JF02671}attP2 | Bloomington | 27521 |
| FBgn0004841 Tkr86C | y[1] v[1]; P{y[+t7.7] v[+t1.8]=TRiP.JF02160}attP2 | Bloomington | 31884 |
| FBgn0005642 wdn | y[1] sc[*] v[1]; P{y[+t7.7] v[+t1.8]=TRiP.GLV21019}attP2 | Bloomington | 35654 |
| FBgn0011288 Snap25 | y[1] v[1]; P{y[+t7.7] v[+t1.8]=TRiP.JF02615}attP2 | Bloomington | 27306 |
| FBgn0014024 Rnp4F | y[1] sc[*] v[1]; P{y[+t7.7] v[+t1.8]=TRiP.GL00383}attP2 | Bloomington | 35457 |
| FBgn0020391 Nrk | y[1] sc[*] v[1]; P{y[+t7.7] v[+t1.8]=TRiP.HMC03875}attP40 | Bloomington | 55184 |
| FBgn0037963 Cad87A | y[1] v[1]; P{y[+t7.7] v[+t1.8]=TRiP.JF03143}attP2 | Bloomington | 28716 |
| FBgn0011582 Dop1R1 | y[1] v[1]; P{y[+t7.7] v[+t1.8]=TRiP.HM04077}attP2 | Bloomington | 31765 |
| FBgn0021764 sdk | y[1] sc[*] v[1]; P{y[+t7.7] v[+t1.8]=TRiP.HMS00292}attP2 | Bloomington | 33412 |
| FBgn0038980 Octbeta1R | y[1] v[1]; P{y[+t7.7] v[+t1.8]=TRiP.HMJ22156}attP40 | Bloomington | 58179 |
| FBgn0039045 CG17119 | y[1] sc[*] v[1]; P{y[+t7.7] v[+t1.8]=TRiP.HMS00213}attP2 | Bloomington | 40823 |
| FBgn0042135 CG18812 | y[1] v[1]; P{y[+t7.7] v[+t1.8]=TRiP.GL01147}attP2 | Bloomington | 42777 |
| FBgn0259483 Mob4 | y[1] sc[*] v[1]; P{y[+t7.7] v[+t1.8]=TRiP.HMC05923}attP40 | Bloomington | 65236 |
| FBgn0259168 mnb | y[1] v[1]; P{y[+t7.7] v[+t1.8]=TRiP.HM05098}attP2 | Bloomington | 28888 |
| FBgn0261613 Oaz | y[1] sc[*] v[1]; P{y[+t7.7] v[+t1.8]=TRiP.GLV21080}attP2 | Bloomington | 35715 |
| FBgn0265194 Trpm | y[1] sc[*] v[1]; P{y[+t7.7] v[+t1.8]=TRiP.HMC02889}attP2/TM3, Sb[1] | Bloomington | 44503 |
| FBgn0266137 Dop1R2 | y[1] sc[*] v[1]; P{y[+t7.7] v[+t1.8]=TRiP.HMC02893}attP2 | Bloomington | 51423 |
| FBgn0267339 p38c | y[1] sc[*] v[1]; P{y[+t7.7] v[+t1.8]=TRiP.HMC05719}attP2 | Bloomington | 64846 |
| FBgn0033459 CG12744 | y[1] sc[*] v[1]; P{y[+t7.7] v[+t1.8]=TRiP.HMC06365}attP40/CyO | Bloomington | 67262 |
| FBgn0027574 CG5815 | w1118; P{GD11776}v22203/TM3 | VDRC | 110272 |
| FBgn0042134 Capr | w[1118]; P{GD5147}v45295 | VDRC | 22203 |

**Supplementary Files**

**Supplementary File 1**. Raw egg counts and p values for figures used in this study. Each tab corresponds to a given gene. Within tab contains the genotype of the RNAi hairpin used, raw egg counts, and p-values.

